# Image-Based Meta- and Mega-Analysis (IBMMA): A Unified Framework for Large-Scale, Multi-Site, Neuroimaging Data Analysis

**DOI:** 10.1101/2025.06.16.657725

**Authors:** Nick Steele, Rajendra A. Morey, Ahmed Hussain, Courtney Russell, Benjamin Suarez-Jimenez, Elena Pozzi, Hadis Jameei, Lianne Schmaal, Ilya M. Veer, Lea Waller, Neda Jahanshad, Sophia I. Thomopoulos, Lauren E. Salminen, Miranda Olff, Jessie L. Frijling, Dick J. Veltman, Saskia B.J. Koch, Laura Nawijn, Mirjam van Zuiden, Li Wang, Ye Zhu, Gen Li, Dan J. Stein, Jonathan Ipser, Yuval Neria, Xi Zhu, Orren Ravid, Sigal Zilcha-Mano, Amit Lazarov, Ashley A. Huggins, Jennifer S. Stevens, Kerry Ressler, Tanja Jovanovic, Sanne J.H. van Rooij, Negar Fani, Sven C. Mueller, Anna R. Hudson, Judith K. Daniels, Anika Sierk, Antje Manthey, Henrik Walter, Nic J.A. van der Wee, Steven J.A. van der Werff, Robert R.J.M. Vermeiren, Christian Schmahl, Julia I. Herzog, Ivan Rektor, Pavel Říha, Milissa L. Kaufman, Lauren A. M. Lebois, Justin T. Baker, Isabelle M. Rosso, Elizabeth A. Olson, Anthony King, Israel Liberzon, Michael Angstadt, Nicholas D. Davenport, Seth G. Disner, Scott R. Sponheim, Thomas Straube, David Hofmann, Guangming Lu, Rongfeng Qi, Xin Wang, Austin Kunch, Hong Xie, Yann Quidé, Wissam El-Hage, Shmuel Lissek, Hannah Berg, Steven E. Bruce, Josh Cisler, Marisa Ross, Ryan J. Herringa, Daniel W. Grupe, Jack B. Nitschke, Richard J. Davidson, Christine Larson, Terri A. deRoon-Cassini, Carissa W. Tomas, Jacklynn M. Fitzgerald, Jeremy Elman, Matthew Panizzon, Carol E. Franz, Michael J. Lyons, William S. Kremen, Brandee Feola, Jennifer U. Blackford, Bunmi O. Olatunji, Geoffrey May, Steven M. Nelson, Evan M. Gordon, Chadi G. Abdallah, Ruth Lanius, Maria Densmore, Jean Théberge, Richard W.J. Neufeld, Paul M. Thompson, Delin Sun

## Abstract

The increasing scale and complexity of neuroimaging datasets aggregated from multiple study sites present substantial analytic challenges, as existing statistical analysis tools struggle to handle missing voxel-data, suffer from limited computational speed and inefficient memory allocation, and are restricted in the types of statistical designs they are able to model. We introduce Image-Based Meta- & Mega-Analysis (IBMMA), a novel software package implemented in R and Python that provides a unified framework for analyzing diverse neuroimaging features, efficiently handles large-scale datasets through parallel processing, offers flexible statistical modeling options, and properly manages missing voxel-data commonly encountered in multi-site studies. IBMMA produced stronger effect sizes and revealed findings in brain regions that traditional software overlooked due to missing voxel-data resulting in gaps in brain coverage. IBMMA has the potential to accelerate discoveries in neuroscience and enhance the clinical utility of neuroimaging findings.

## Introduction

As the field of neuroimaging has matured, there has been a growing recognition of the limitations inherent in single-cohort studies that suffer from small sample sizes coupled with high-dimensional data that reduce statistical power, site-specific biases in data collection and processing, limited demographic diversity that reduces generalizability, and insufficient representation of illness subtypes or rare conditions. To address these challenges, the neuroimaging community has increasingly embraced big data approaches by aggregating datasets from multiple study sites to enhance statistical power, improve reproducibility, and capture the rich heterogeneity of brain function and structure sourced from diverse samples. This paradigm shift has been spurred by multi-site studies and neuroimaging consortia, which offer the potential for more robust, generalizable findings that can significantly advance our understanding of the brain.

To facilitate the complex task of preprocessing and analyzing neuroimaging big data, the scientific community has developed a suite of standardized pipelines and tools. Open-source solutions such as fMRIPrep (Esteban et al., 2019), CONN toolbox (Whitfield-Gabrieli & Nieto-Castanon, 2012), Analysis of Functional NeuroImages (AFNI) (Cox, 1996), and FMRIB Software Library (FSL) (Jenkinson et al., 2012) provide comprehensive workflows for data preprocessing and/or analysis. Building on this foundation, the ENIGMA (Enhancing NeuroImaging Genetics through Meta-Analysis) Consortium introduced HALFpipe (Waller et al., 2022), a sophisticated containerized pipeline that extends fMRIPrep’s workflow and incorporates additional preprocessing steps, quality assessment tools, and feature extraction capabilities. However, limitations not addressed by HALFpipe, such as modeling complex site effects and unified integration of meta- and mega-analysis, remain unresolved.

We have developed Image-Based Meta- & Mega-Analysis (IBMMA), a software package tailored for analyzing neuroimaging datasets aggregated from multiple study sites. IBMMA unifies the analysis of diverse neuroimaging features, including voxelwise, vertexwise, and connectome-wide measures, while efficiently managing large-scale datasets by leveraging parallel processing capabilities common in modern high performance clusters (HPC). The IBMMA package provides flexible statistical modeling options, model specification in simple text format files, robust handling of missing data by dynamically varying the degrees of freedom for each neuroimaging feature (e.g., voxel or vertex), and generates interpretable outputs that preserve the original data structure. These features enable comprehensive analysis of complex multi-site neuroimaging studies not available with other neuroimaging software (**Table 1**).

**Table 1:**
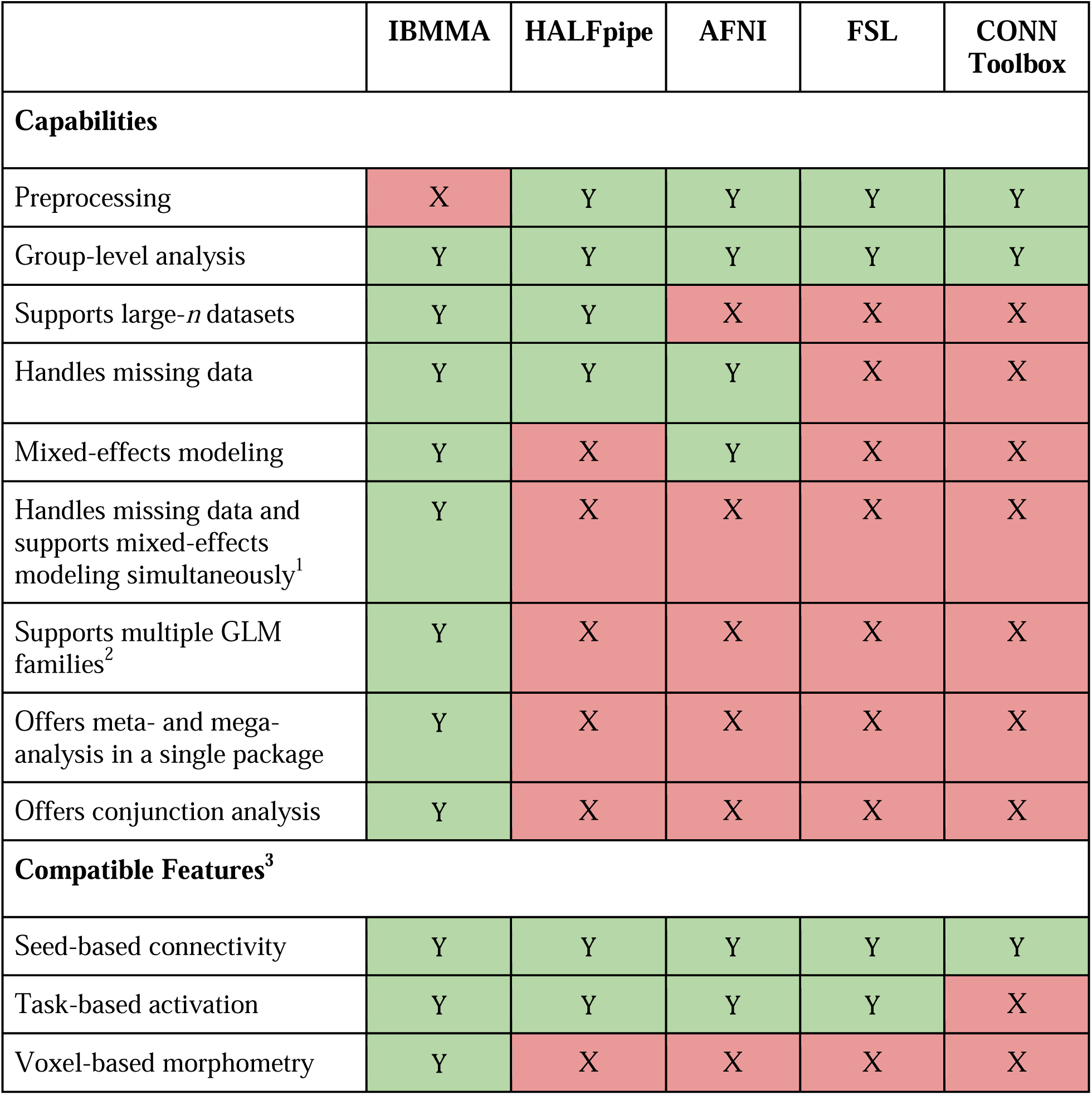
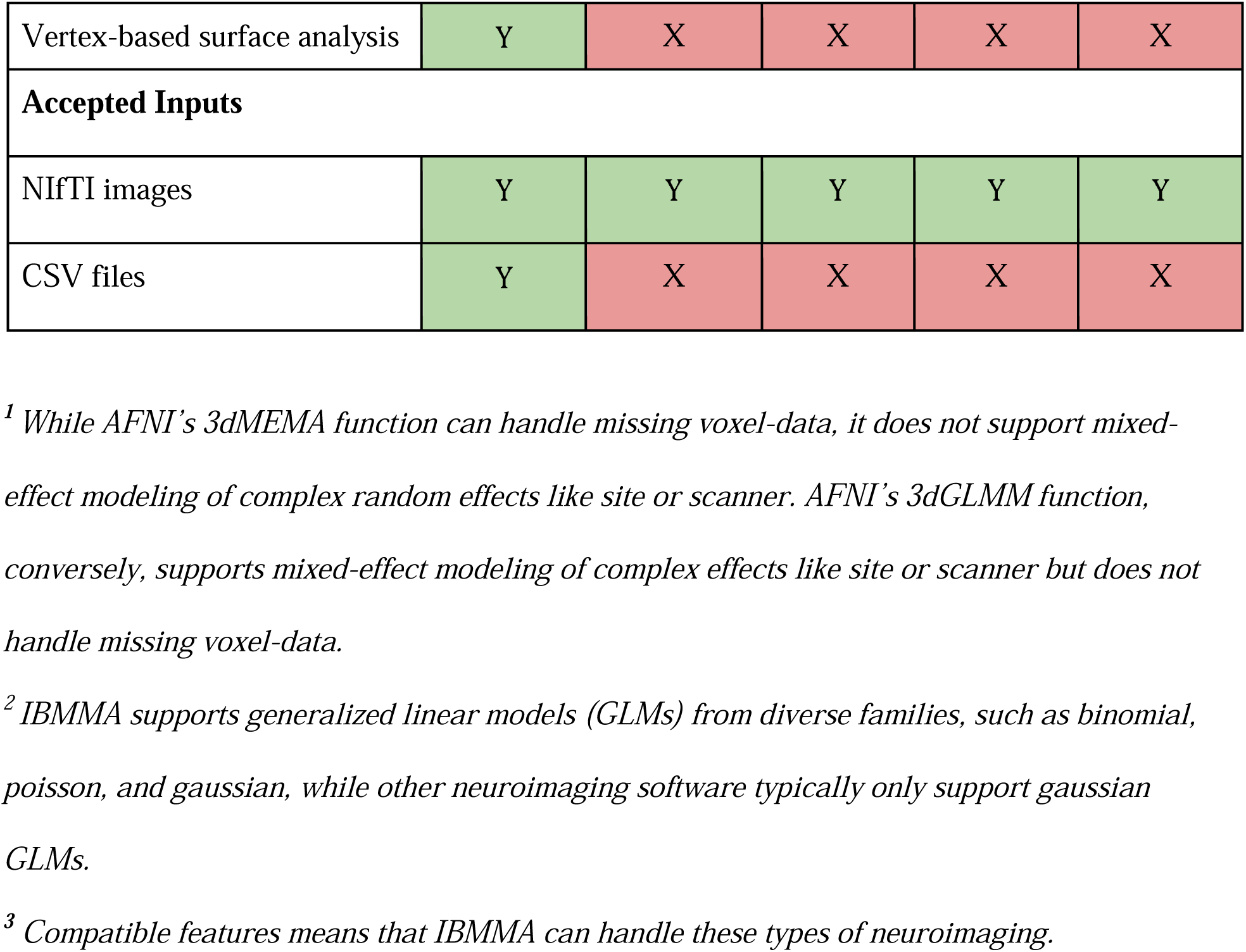
Comparison between IBMMA and other neuroimaging software. ^1^.

|  | IBMMA | HALFpipe | AFNI | FSL | CONN<br>Toolbox |
| --- | --- | --- | --- | --- | --- |
| **Capabilities** |  |  |  |  |  |
| Preprocessing | X | Y | Y | Y | Y |
| Group-level analysis | Y | Y | Y | Y | Y |
| Supports large-*n* datasets | Y | Y | X | X | X |
| Handles missing data | Y | Y | Y | X | X |
| Mixed-effects modeling | Y | X | Y | X | X |
| Handles missing data and supports mixed-effects modeling simultaneously<sup>1</sup> | Y | X | X | X | X |
| Supports multiple GLM families<sup>2</sup> | Y | X | X | X | X |
| Offers meta- and mega-analysis in a single package | Y | X | X | X | X |
| Offers conjunction analysis | Y | X | X | X | X |
| **Compatible Features<sup>3</sup>** |  |  |  |  |  |
| Seed-based connectivity | Y | Y | Y | Y | Y |
| Task-based activation | Y | Y | Y | Y | X |
| Voxel-based morphometry | Y | X | X | X | X |
| Vertex-based surface analysis | Y | X | X | X | X |
| **Accepted Inputs** |  |  |  |  |  |
| NIfTI images | Y | Y | Y | Y | Y |
| CSV files | Y | X | X | X | X |

<sup>1</sup> While AFNI's 3dMEMA function can handle missing voxel-data, it does not support mixed-effect modeling of complex random effects like site or scanner. AFNI's 3dGLMM function, conversely, supports mixed-effect modeling of complex effects like site or scanner but does not handle missing voxel-data.
<sup>2</sup> IBMMA supports generalized linear models (GLMs) from diverse families, such as binomial, poisson, and gaussian, while other neuroimaging software typically only support gaussian GLMs.
<sup>3</sup> Compatible features means that IBMMA can handle these types of neuroimaging.

Large-scale neuroimaging consortia have predominantly employed two approaches to analyze data aggregated from multiple sites: meta-analysis and mega-analysis. Meta-analysis typically involves each site independently generating results in the form of sample means, sample variances, and effect size estimates for each model being tested, which are then used to calculate a total effect size estimate across sites (Radua et al., 2020). By contrast, mega-analysis involves pooling individual participant data from multiple studies or sites into a single sample, and then applying statistical modeling to the pooled sample (Sun et al., 2022; Thompson et al., 2020). However, traditional software packages, particularly for neuroimaging data, have treated these approaches separately, requiring researchers to use different tools that reside in different packages for meta- and mega-analyses.

Another significant challenge in fMRI analyses is missing data that often presents near cortical boundaries, the medial temporal lobe, and the ventral prefrontal cortex (Mulugeta et al., 2017; Vaden et al., 2012). These missing values most often stem from signal dropout due to magnetic susceptibility artifacts near air-tissue interfaces, head motion during scanning, and inconsistent scanning protocols and spatial normalization procedures across sites. The challenge becomes particularly acute in multi-site studies, where missing values often vary systematically by site. This can result in substantial data loss, effectively reducing the brain voxels/regions being analyzed as the sample size increases Traditional analysis approaches that simply omit voxels or vertices with incomplete observations can significantly compromise the quality of the results. This may lead to both inflated Type II (false negative) error rates from excluded voxels or vertices, and inflated Type I (false positive) error rates, as multiple comparison corrections are applied to fewer voxels or vertices than the total number in the brain.

The complexity of statistical modeling presents another major challenge in large-scale neuroimaging analyses. Traditional software tools support limited types of statistical models, typically restricted to gaussian family fixed-effect generalized linear models (GLMs) for group-level analysis. This restricts researchers from applying more flexible statistical approaches, such as mixed-effects models, which are essential for properly accounting for variability across subjects, scanner types, and study sites in multi-site studies (Thompson et al., 2020). For instance, while FSL’s FLAME uses linear mixed models for Bayesian estimation of subject-level variability, it cannot model more complex random effects such as scanner or site. AFNI’s 3dLMEr and 3dGLMM functions calls the *lme4* R package and can be used to model scanner- or site-related variability, but it was not originally designed for large-*n* datasets and does not properly handle missing neuroimaging data. Most specialized neuroimaging tools (e.g., AFNI, SPM, FSL, FreeSurfer) struggle with datasets containing thousands of subjects, as the memory requirements for processing such large-scale data often exceed standard computer capabilities, leading to computational failures.

Additionally, general-purpose statistical packages (e.g. SAS, SPSS, R) are designed for univariate analyses but struggle with mass-univariate neuroimaging data due to non-vectorizable operations across features. Big Linear Mixed Models (BLMM) is a Python tool designed for analyzing mass-univariate linear mixed models in large-scale neuroimaging datasets (Maullin-Sapey & Nichols, 2022), addressing challenges posed by increasing data complexity and size. It leverages vectorization for improved performance across multiple voxels and uniquely accounts for "voxelwise missingness". However, BLMM is limited to mega-analysis of voxelwise images using linear mixed models and does not support robust multiple comparison corrections.

We demonstrate the capabilities of IBMMA to address these limitations by applying it to resting-state fMRI data from the ENIGMA-PTSD working group. This analysis demonstrates the abilities of IBMMA to conduct meta- and mega-analyses on multi-site functional neuroimaging data. We compare the results generated by IBMMA to those generated by FSL and highlight the advantages and disadvantages of each approach.

## Methods

### IBMMA Workflow

#### Mega-Analysis

IBMMA processes data through several sequential steps. First, *Data Preparation* identifies relevant files based on user specifications. The *Data Masking* step applies subject-specific whole-brain or gray matter masks to exclude irrelevant features, such as voxels outside the brain, and a standardized (non-subject-specific) brain mask to constrain the final output. All masks must match the dimensionality of the imaging data. IBMMA includes predefined standard space whole-brain masks compatible with commonly used neuroimaging tools, including a 97×115×97 voxel mask (compatible with HALFpipe) and a 91×109×91 voxel mask (compatible with FSL). Alternatively, users can provide custom masks to suit their analysis needs.

As shown in **Fig. 1**, the *Data Flattening* step converts all data types to one-dimensional vector format for uniform processing. *Data Segmentation and Stacking* then organizes the flattened data into subject-by-feature matrices and divides them into memory-compatible segments. The number of data segments can be adjusted by the user based on available memory resources to optimize parallel processing efficiency.

**Figure 1.**
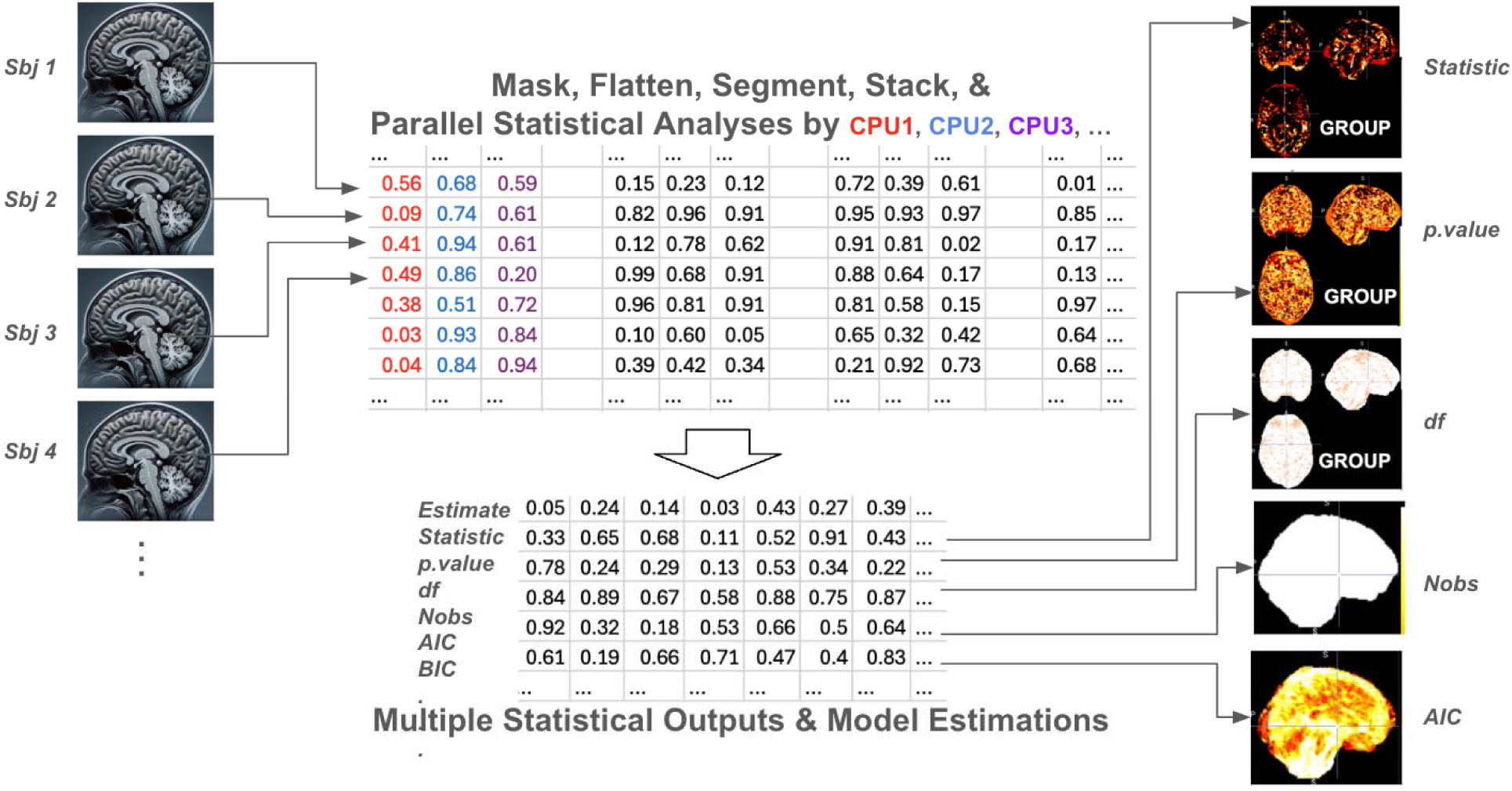
Schematic of IBMMA Workflow. Neuroimaging data from multiple subjects are flattened and segmented into memory-compatible subject-by-feature matrices. Statistical models are applied in parallel across features, and the resulting outputs are extracted and reconstructed to match the original data dimensions.

The *Statistical Analysis* step executes parallel computations of each neuroimaging feature using specified R or Python scripts. IBMMA currently supports regression models using functions from the *stats* and *lme4* R packages, including *lm()* for linear models, *glm()* for generalized linear models, *lmer()* for linear mixed-effects models, and *glmer()* for generalized linear mixed-effects models. Finally, *Results Compilation* extracts statistical outputs using the *broom* or *broom.mixed* R packages, merges results from all segments, and reconstructs them to match the original data dimensions. IBMMA generates multiple statistical outputs, including coefficient statistics (effect size estimates, standard errors, p-values) and model fit measures (R-squared, AIC, BIC).

For multiple comparison correction, IBMMA is able to perform false discovery rate (FDR) correction and probabilistic threshold-free cluster enhancement (pTFCE) (Spisák et al., 2019) with family-wise error rate correction. The pTFCE approach, particularly valuable for voxelwise images, addresses limitations of traditional cluster inference methods by offering enhanced robustness to cluster topology, strict control over false discovery, and improved sensitivity compared to conventional TFCE (Smith & Nichols, 2009). This enables straightforward multiple comparison correction and avoids time-consuming permutation testing by using Bayesian probability calculations.

#### Meta-Analysis

IBMMA performs meta-analysis by applying the statistical modeling pipeline described above to each study site independently. This site-wise analysis generates complete summary statistics for each neuroimaging feature, including effect size estimates, standard errors, test statistics, and *p*-values, along with sample size information for meta-analytic weighting. Site-specific summary statistics are then aggregated to calculate meta-analytic results using NiMARE (Neuroimaging Meta-Analysis Research Environment) (Salo et al., 2023). This calculation is performed using either Stouffer’s method (default) or Fisher’s method. Stouffer’s method combines *Z*-scores across studies, optionally weighting them by sample size, and divides the weighted sum by the square root of the sum of squared weights (or the square root of the number of studies in the unweighted case). The resulting combined *Z*-score is compared to the standard normal distribution to assess statistical significance. Fisher’s method, by contrast, combines *p*-values by summing the negative logarithms of the *p*-values, yielding a chi-square statistic whose significance is assessed against the chi-square distribution.

#### Conjunction Analysis

IBMMA supports multiple forms of conjunction analysis, which allows for direct comparison of meta-analytic and mega-analytic results to identify consistent findings across both analytic strategies. Conjunction analysis can be used more broadly to compare results from different preprocessing strategies, original and replication datasets, or outputs from different statistical models or software packages. One method of conjunction analysis offered by IBMMA is overlap (logical AND) conjunction, which identifies the intersection of neuroimaging features that are statistically significant across inputs.

Other conjunction analysis methods available include the minimum or strict conjunction approach, which takes the minimum test statistic across inputs and only identifies effects that are significant in all inputs. The conjunction null method (Nichols et al., 2005) tests whether at least one effect is null and only identifies regions where all effects are statistically significant, providing a more rigorous alternative to simple overlap. The global or lenient conjunction method is based on averaging test statistics to detect consistent effects across inputs, even if not all are individually significant. Finally, the probabilistic approach estimates the joint probability of effects across inputs using a Bayesian-inspired framework, allowing for more graded and interpretable conclusions than binary conjunctions.

### Running IBMMA

#### Installation

IBMMA is an open-source software that can be downloaded from the developer’s GitHub (https://github.com/sundelinustc/IBMMA). The current version was developed and tested using Python 3.11.9 and R 4.2.2. Before running IBMMA, users must install required dependencies. Software configuration is managed via the *para_path.xlsx* file, which specifies: (1) the file path to a clinical or behavioral data file, (2) the file path to the directory containing all neuroimaging data organized into site-specific subdirectories, (3) a filename pattern that IBMMA uses to identify relevant files within the neuroimaging data folder, (4) a list of predictor variables for statistical analysis, and (5) the statistical models to be executed.

#### Data Preparation

IBMMA requires a structured clinical or behavioral dataset containing subject-specific variables for analysis. When constructing an IBMMA-compatible clinical or behavioral data file, users must assign each subject with a *fID,* which combines the *Site ID* and *Subject ID* to ensure that each subject has a unique identifier (**Fig. 4A, right**). While *Subject ID*s may be shared between different sites, *fID*s remain unique for every subject. For example, if both *Site1* and *Site2* use four-digit *Subject ID*s (e.g., *1234*), IBMMA expects corresponding *fID*s such as *Site1_1234* and *Site2_1234*.

Users’ neuroimaging data is expected to be organized in site-specific subdirectories with file names beginning with the *Subject ID* (**Fig. 2A, left**). To ensure proper file recognition, all neuroimaging files must follow a standardized naming convention that appends a consistent pattern to the *Subject ID*. Users must specify this naming pattern in the *data_pattern* tab of the *para_path.xlsx* file. IBMMA then recursively searches all subdirectories within the designated data directory (defined in the *data_path* tab) and selects files that match the specified pattern. Finally, users define predictor variables and statistical models in the *para_path.xlsx* file under the *predictors* and *models* tab, respectively. Statistical models should be formatted following the standard syntax for the chosen function (**Fig 2B**).

**Figure 2.**
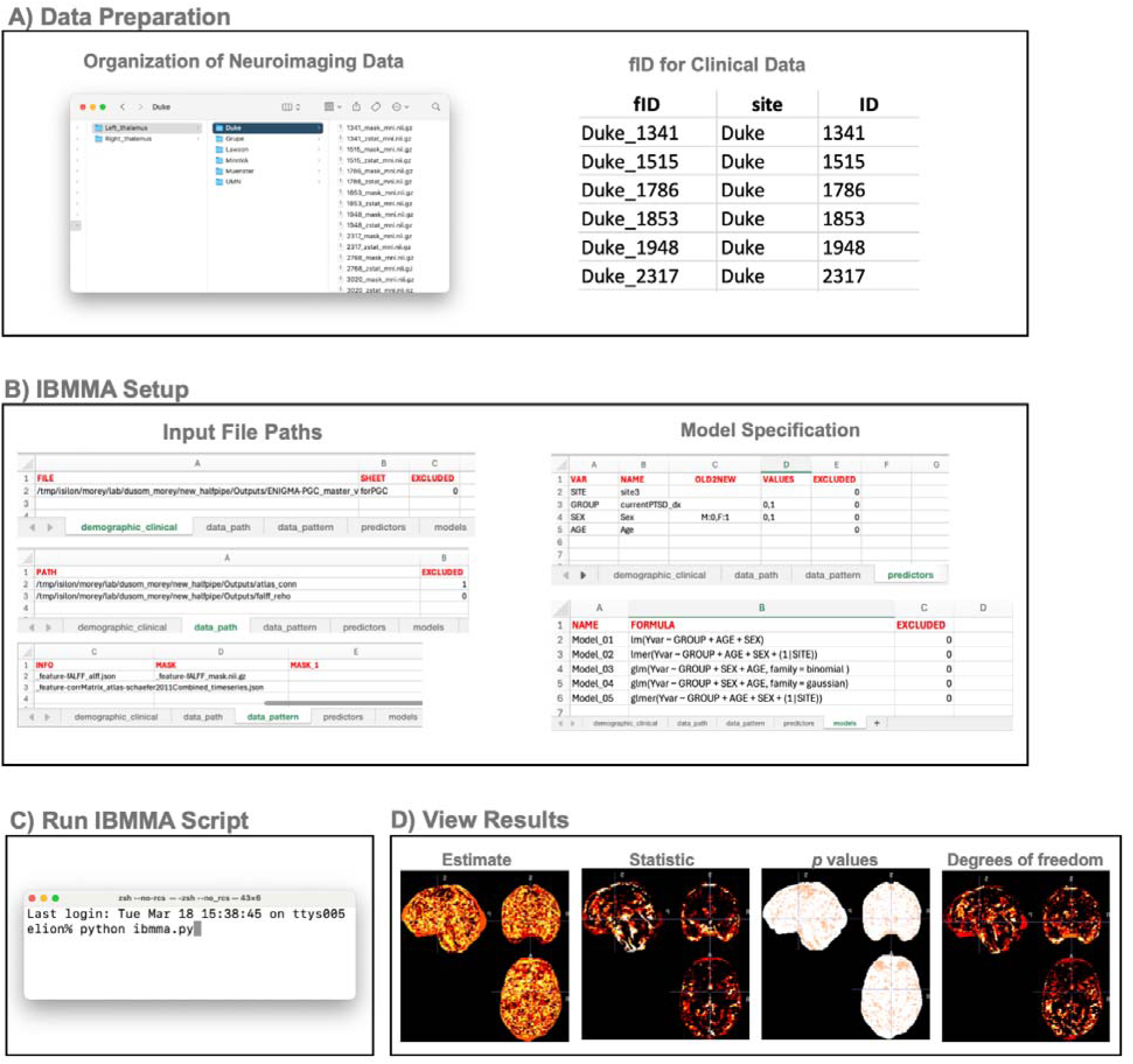
Steps to Run IBMMA. **A)** IBMMA expects neuroimaging data to be organized in site-specific subdirectories. Within each subdirectory, neuroimaging files should be named starting with the subject ID followed by a naming pattern that is consistent across all files. Within the clinical data file, IBMMA expects a fID column whose elements are composed of site ID and subject ID. **B)** IBMMA requires the user to first input clinical and neuroimaging data file paths and model specifications in the para_path.xlsx file. **C)** IBMMA can be run from the command line, either locally or via a HPC, by running the ibmma.py script. **D)** Statistical results will be automatically generated once the script is done running and can be viewed using any neuroimaging viewer (e.g., FSLeyes, freeview, MRIcron). An HTML webpage is also generated to view results.

### Application to ENIGMA-PTSD Dataset

To demonstrate IBMMA’s capabilities, we applied it to resting-state fMRI data from the ENIGMA PTSD working group. This dataset includes 3,193 participants from 29 study sites. After removing participants missing essential clinical data, such as age, sex, or PTSD diagnosis, 2,282 participants remained for analysis with IBMMA (**Table S1**).

#### Data Preprocessing

Data were preprocessed using HALFpipe, which included removal of the first four volumes, grand mean scaling, motion correction, highpass temporal filtering, denoising with ICA-AROMA (Pruim et al., 2015), and spatial normalization.

#### Statistical Model

We implemented a linear mixed-effects model to examine the associations between age, sex, and PTSD diagnosis with left thalamus resting-state functional connectivity (RSFC). The model was specified as (1) Brain ∼ Age + Sex + Dx + (1|Site), where ’Brain’ represents the voxelwise neuroimaging measure, ’Dx’ is the diagnostic status (0 = control, 1 = case), and ’Site’ was treated as a random effect to account for site-specific variability (intercept only).

#### IBMMA Execution

IBMMA was executed on the Duke-UNC BIAC high-performance computing cluster using a single node with 48 cores per CPU. The job requested 12-core linear binding, two processing cores, and 100GB of both virtual and physical memory to accommodate the computational demands of large-scale neuroimaging analyses. IBMMA generated whole-brain statistical maps, which were then visualized to identify brain regions showing associations with diagnosis, age, and sex. Statistical results were thresholded at *Z* ≥ 3.1 without correction for multiple comparisons, as the focus was on facilitating qualitative visualization and comparisons rather than assessing statistical significance.

#### Comparisons with FSL

IBMMA results were compared to FSL outputs generated by analyzing resting-state fMRI data consisting of 397 participants from 6 study sites (**Table S2**). Only a subset of our sample was used for this comparison due to the aforementioned challenges with processing a large-scale dataset using traditional neuroimaging software. Data were preprocessed using FSL (Jenkinson et al., 2012), including removal of the first four volumes, slice-timing correction, motion correction, highpass temporal filtering, spatial normalization, and noise reduction using ICA-AROMA (Pruim et al., 2015). First-level RSFC analysis was conducted in FSL using the left thalamus as a seed for each participant. Seed-whole brain voxelwise *Z*-statistic maps were fed into IBMMA and FSL for group-level analysis. A GLM was used to examine associations with age, sex, and diagnosis. The model was specified a s (2) Brain ∼ Age + Sex + Dx. Overlap conjunction analysis of the resulting *Z*-statistic images was used to assess the degree of agreement among the software packages, enabling direct comparison of their relative sensitivity and statistical power.

### Simulated Data Analysis

A simulated neuroimaging dataset with varying levels of missing data was used to evaluate the performance of IBMMA. The simulated dataset was designed to mirror realistic conditions encountered in large-scale neuroimaging studies where data missingness is common due to technical artifacts, motion, or quality control procedures. We simulated 1,000 brains as 50×50×50 voxel arrays, totaling 125,000 voxels. Voxel values were drawn from a standard normal distribution (μ = 0, σ = 1).

To establish a ground truth signal, we embedded a spherical region of interest with a radius of 4 voxels, totaling 251 voxels. A binary diagnostic covariate with 50% prevalence was used to represent case-control study designs commonly used in clinical neuroimaging research. Within the signal region, we applied an effect size of Cohen’s *d* = 0.2 by adding a value of 0.2 to signal voxels for case subjects. This effect size was chosen to reflect the modest but clinically meaningful effects typically observed in neuroimaging studies of psychiatric conditions (e.g, Schmaal et al., 2020).

The impact of missing data on IBMMA’s performance was evaluated by introducing missing data in 5% increments, ranging from 0% to 100% of subjects. For each level of missing data, receiver operating characteristic (ROC) curves were generated, and two performance metrics were computed: area under the curve (AUC) and sensitivity at a 5% false positive rate (FPR). Additionally, IBMMA was compared to complete-case analysis, where only voxels with no missing data are analyzed, using a missing completely at random (MCAR) mechanism to introduce missing data, which is reported in our *Supplementary Materials*.

## Results

### Mega- and Meta-Analysis

IBMMA successfully performed a mega-analysis by analyzing voxelwise RSFC data from 2,282 participants across 29 study sites using a linear mixed-effects model (model 1). Total run time, CPU time, and maximum virtual memory usage for each processing step are reported in **Fig 3A**. Statistical results were generated within the *Results* directory, including whole-brain maps of statistical estimates, *Z*-statistics, pTFCE *Z*-statistics, *p*-values, FDR-corrected *p*-values, and standard errors. Whole-brain maps of model fit measures were additionally generated, including number of observations, degrees of freedom, residual degrees of freedom, Akaike information criterion, Bayesian information criterion, log likelihood, restricted maximum likelihood, and sigma (**Fig. 3B**). IBMMA automatically produced an HTML webpage summarizing key results (**Fig. 3C**). This included tables with demographic and clinical statistical results, site-specific demographic and clinical information, a table of all significant voxel clusters, and interactive brain maps to view results.

**Figure 3.**
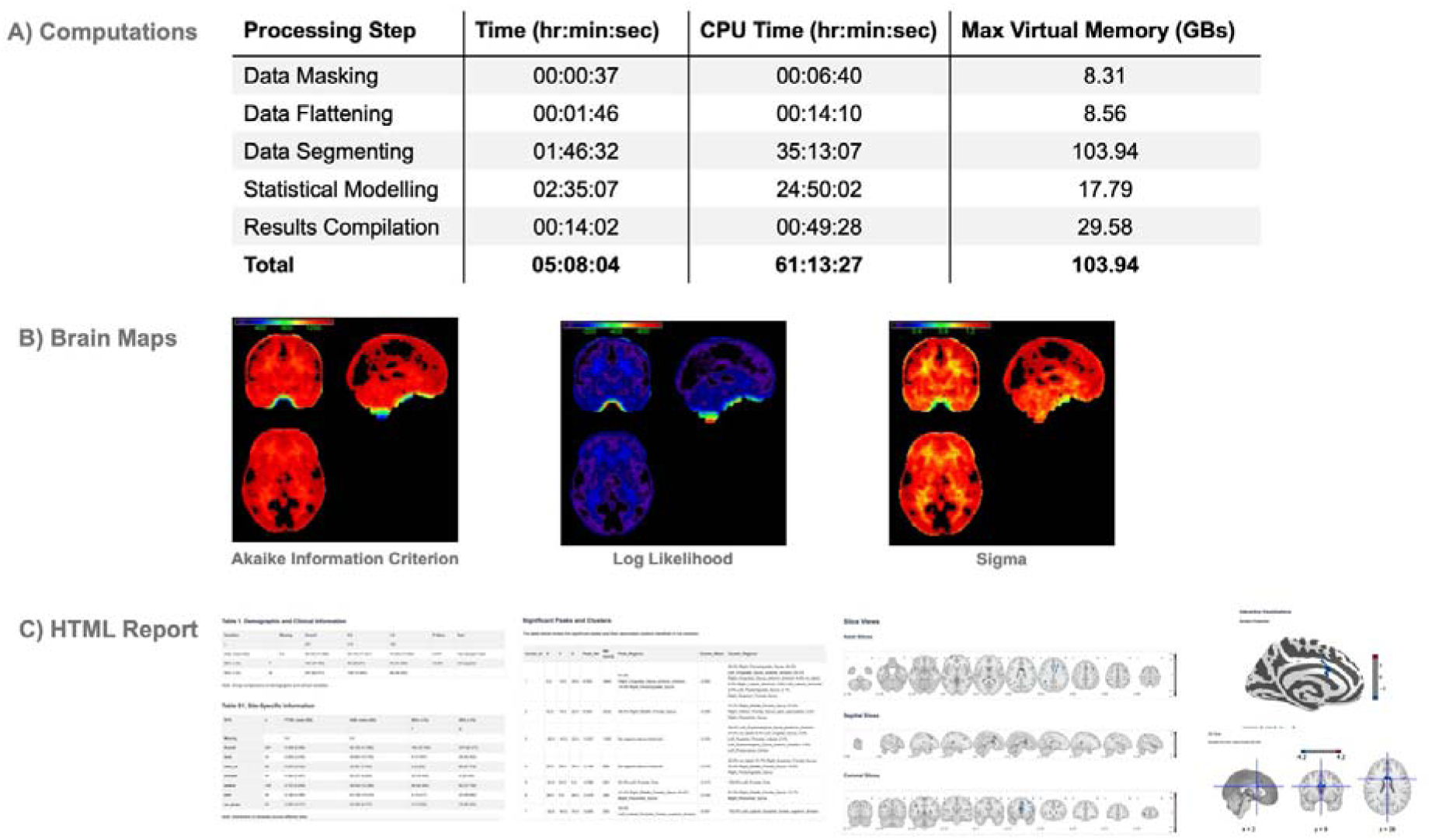
IBMMA Outputs. **A)** Total run time, CPU time, and maximum virtual memory usage of each processing step of IBMMA’s mega-analysis using a sample of n = 2,611. **B)** IBMMA generated whole-brain maps of various statistical and model fit measures. **C)** An automatically generated HTML webpage provided result tables and visualizations of neuroimaging findings. All tables were also saved as CSV files, and all significant voxel clusters were saved as png images.

Results for the meta-analysis were generated in the *Results* directory, which contained an HTML webpage with result summaries and a whole-brain t-statistic map. Overall, meta-analysis results had fewer voxels surviving the *Z* ≥ 3.1 threshold but were nonetheless consistent with the mega-analysis results.

To assess the impact of missing data on model performance across the brain, we conducted a voxelwise analysis of subject missingness and evaluated the sensitivity of the IBMMA method under varying levels of data loss using simulated data. **Fig. 4A** displays a brain map of the percentage of missing subjects at each voxel, computed from the full ENIGMA-PTSD dataset (*n* = 3,193). This brain map highlights regions with higher susceptibility to data loss due to factors such as variable brain coverage or susceptibility artifacts.

**Figure 4.**
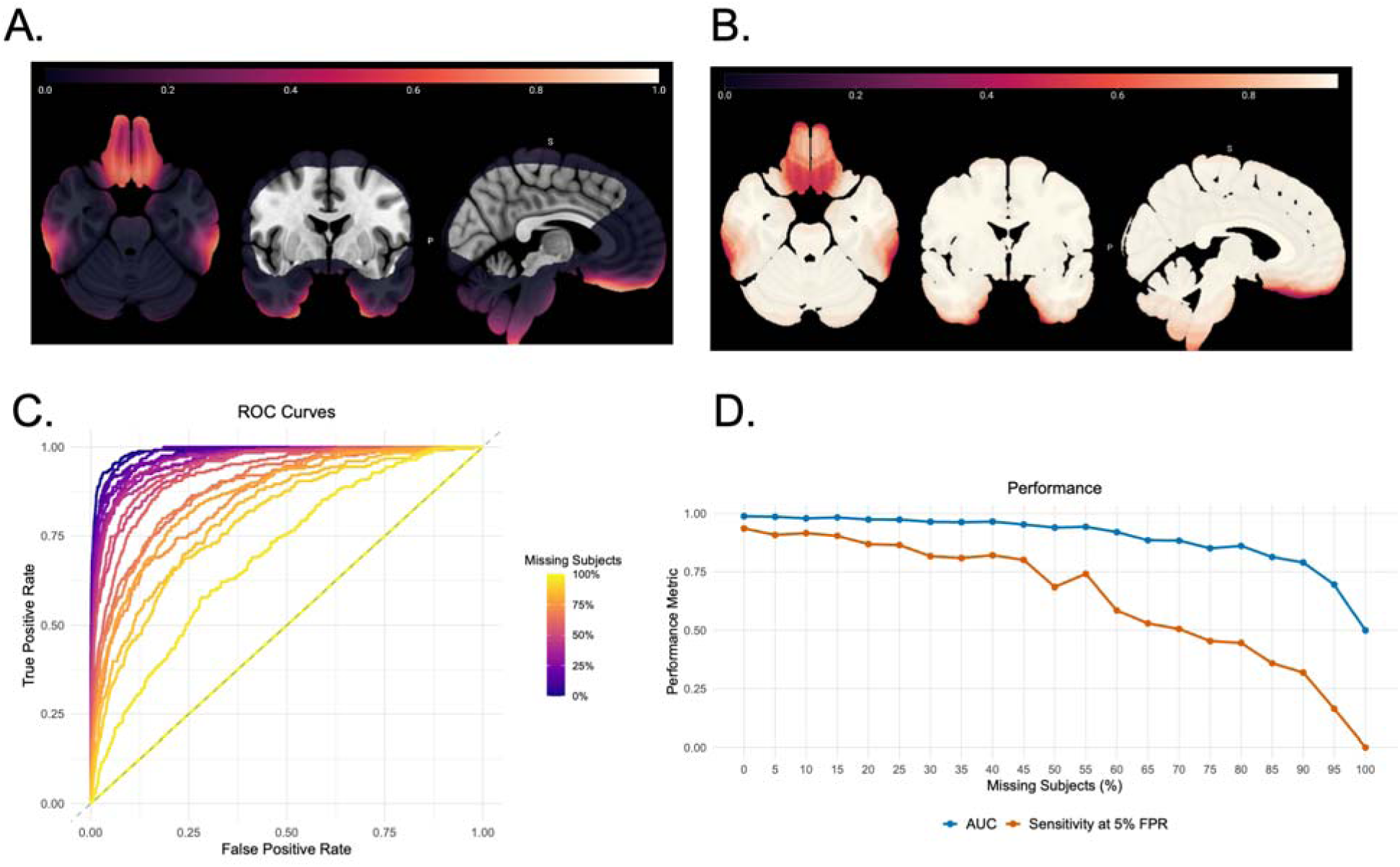
**A)** Brain maps displaying the proportion of subjects with missing data at each voxel of the brain in the ENIGMA-PTSD dataset (n = 3,193). **B)** The sensitivity of IBMMA at a 5% false positive rate (FPR) calculated from a simulated dataset given the observed missing rates at each voxel of the brain. **C)** Receiver operator characteristic (ROC) curves at different levels of subject missingness in increments of 5%. **D)** Performance of IBMMA, measured by area under the curve (AUC) and sensitivity at a 5% FPR, across varying levels of subject missingness.

We then evaluated IBMMA’s performance across increasing levels of missing subjects. For each 5% increment of missingness (from 0% to 100%), we generated a receiver operating characteristic (ROC) curve, computing the area under the curve (AUC) and sensitivity at a 5% FPR. These results are summarized in **Fig. 4C**, where ROC curves illustrate the decline in performance as missing data increases, and **Fig. 4D**, where line plots show the performance of IBMMA measured by AUC and sensitivity at a 5% FPR across increasing levels of missing data. Using the voxelwise missingness percentages from **Fig. 4A**, we assigned to each voxel the sensitivity of IBMMA at a 5% FPR corresponding to its level of missing data. The resulting brain map, shown in **Fig. 4B**, visualizes the expected sensitivity of IBMMA across the brain given the observed missingness profile.

### Mega-Analysis Comparison with FSL

IBMMA produced statistical results in brain regions which lacked coverage in the FSL analysis. Large dorsal and ventral brain areas were excluded by FSL from its analysis and results due to missing voxel-data for one or more participants in those regions. IBMMA’s voxelwise analysis did not exclude any voxels because IBMMA only included participants for whom voxel-data was present for a given voxel (**Fig 5A**).

**Figure 5.**
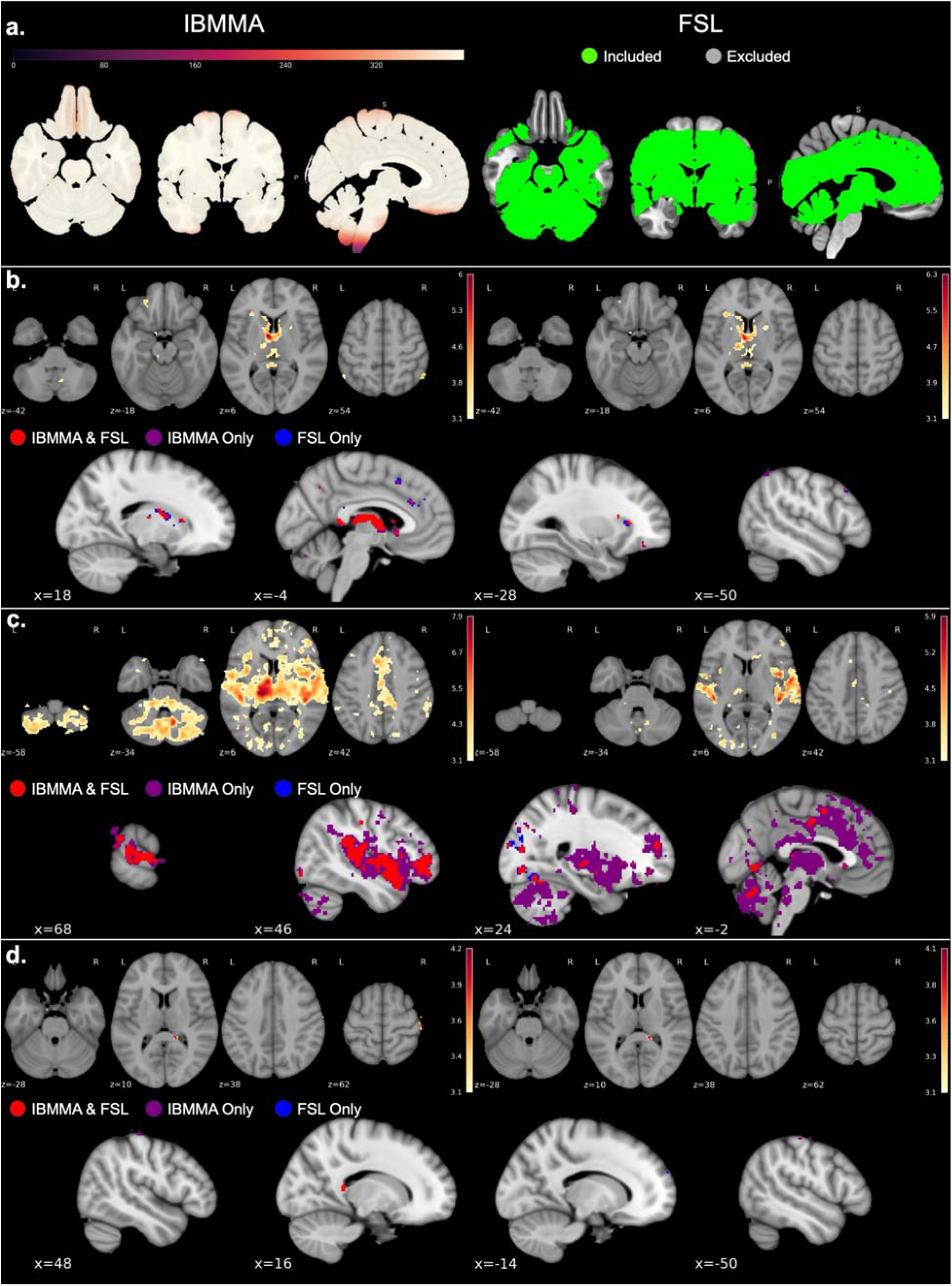
Seed-based functional connectivity results for the left thalamus using IBMMA, FSL, and conjunction analyses. **A)** IBMMA accounts for missing voxel-data by only using subjects with non-missing data for a given voxel, while FSL only analyzes voxels with complete data. Left: Number of participants with non-missing voxel-data used by IBMMA for each voxel. Right: The colored regions represent voxels that were included in the FSL analysis, while non-colored regions were excluded due to one or more participants having missing voxel-data. **B, C, D)** Significant associations found by IBMMA (left) and FSL (right). Z-statistic maps are thresholded at Z ≥ 3.1 (uncorrected). In the conjunction analysis panels (bottom), red voxels indicate regions identified as significant by both IBMMA and FSL, while purple (IBMMA only) and blue (FSL only) voxels indicate areas of disagreement between the two software. **B)** Significant negative associations with age. **C)** Significant associations with sex (female > male). **D)** Significant association with PTSD diagnosis (cases > controls).

IBMMA and FSL analyses revealed associations of left thalamus RSFC with age, sex, and diagnosis (*Z* ≥ 3.1, uncorrected; model 2). Results between the two software were largely consistent with each other (**Fig. 5**). To test the degree of similarity in results, the IBMMA and FSL whole-brain *Z*-statistic maps were masked using a brain mask that included only voxels present in both analyses, and the correlation between the results was computed. The IBMMA and FSL *Z*-statistic maps for associations with age, sex, and diagnosis had a Pearson’s correlation of *r* = 0.97, *r* = 0.76, and *r* = 0.92, respectively. Overlap conjunction analysis was performed using IBMMA’s conjunction analysis module to visualize similarities and differences in results between IBMMA and FSL. **Fig. 5** shows where the two software agreed (red voxels) and disagreed (purple and blue voxels).

## Discussion

We developed Image-Based Meta- & Mega-Analysis (IBMMA), which is a novel software package designed for efficient analysis of large-scale, multi-site neuroimaging data. Results of applying the IBMMA package to resting-state fMRI data from ENIGMA-PTSD are presented as a test case. Our findings support the utility and technical advances of IBMMA in addressing key challenges when analyzing neuroimaging big data. We demonstrated that IBMMA addressed major challenges, including (1) being capable of mega-analyzing large datasets of several thousand samples in a reasonable timeframe, (2) analyses of each voxel being performed in the participants with available voxel-data, whereas participants with missing voxel-data are omitted, (3) performing mixed-effects modeling to support multi-site data analysis, (4) performing both meta-analysis and mega-analysis in a single package and performing conjunction analysis, and (5) automatically producing statistical results tables and figures, allowing efficient and consistent reporting of results.

Missing voxelwise data poses a significant challenge in neuroimaging analyses. Many existing software packages, including FSL, require complete data across all participants for a given voxel to include it in the analysis. As a result, entire brain regions may be excluded from statistical testing if even only a single participant lacks data at those voxels. This restriction reduces spatial coverage and may obscure meaningful group-level effects, especially in regions prone to signal dropout or partial brain coverage. By using only available voxel-data at each voxel, IBMMA ensured that all brain voxels/regions could be analyzed. This facilitated IBMMA in uncovering differences in left thalamus RSFC with the dorsal postcentral gyrus in PTSD, which FSL did not detect due to the latter region lacking voxel-data in one or more participants.

However, the most dorsal and ventral regions of the brain had notably smaller sample sizes due to a higher proportion of missing voxel-data. IBMMA tracks and reports the number of observations contributing to each analysis, allowing researchers to assess the reliability of results, particularly in regions with smaller sample sizes. Using simulated data, we showed that IBMMA displays high performance, even in brain regions with 40% or more missing data. With no missing data, IBMMA achieved an AUC of 0.989 and sensitivity at 5% FPR of 0.936. Performance remained robust, for instance, at 40% subject missingness, with an AUC value above 0.96 and sensitivity above 0.80.

The flexibility of IBMMA’s statistical modeling options facilitated the implementation of a linear mixed-effects model, which enabled IBMMA to account for complex site effects by modeling site as a random intercept. This approach provides a more nuanced and potentially more accurate analysis than traditional fixed-effects models commonly used in neuroimaging studies (Chen et al., 2018). Modeling site as a random effect improves generalizability by extending inferences to sites beyond the specific sites included in the dataset, making findings more applicable to a broader population (Hall & Rosenthal, 2018). This strengthens the interpretation of results as neural correlates for a given disorder or behavior because it helps segregate biological effects from site-specific variability. Future efforts will expand on this capability by incorporating other GLM families, such as binomial models for logistic regression, to address a wider range of research questions. By contrast, many existing tools lack the ability to implement such models, restricting the types of analyses that can be conducted. By offering flexible modeling options, IBMMA strengthens as well as broadens the scope and scale of statistical inference in neuroimaging research.

Overall, our findings highlight IBMMA as a powerful alternative to traditional neuroimaging software, particularly for large multi-site neuroimaging data where missing voxel-data is not uncommon. IBMMA successfully processed and analyzed a large, complex dataset from multiple study sites, demonstrating its ability to handle the computational demands of neuroimaging big data. Software parallelization within IBMMA on a multi-CPU operating system resulted in significant reductions in computation time, addressing a key challenge in the analysis of large-scale neuroimaging studies. IBMMA effectively managed missing voxel-data and accounted for site-specific variance, two issues of paramount importance in multi-site neuroimaging research that have heretofore been challenging (Thompson et al., 2020). By leveraging IBMMA’s capabilities, large-scale consortia can effectively address site-effects when integrating data from diverse samples to discover robust, generalizable patterns of brain activity that are associated with clinical and demographic features.

While IBMMA was developed for neuroimaging applications, its core architecture is designed to handle any high-dimensional data that can be represented as subject-by-feature matrices. The framework accepts CSV-formatted input data and applies statistical models to each feature independently, making it applicable to genomic datasets of gene expression levels or single nucleotide polymorphisms, multi-site clinical data with high-dimensional biomarkers, ecological datasets with spatial or temporal features, or any domain requiring feature-wise statistical modeling.

### Limitations and Future Directions

Despite IBMMA capabilities and the present demonstration of its strengths, several limitations should be noted. Our analysis focused on resting-state fMRI data; future applications of IBMMA will explore other imaging modalities. From a methodological perspective, while IBMMA effectively handled missing voxel-data, future versions will incorporate more sophisticated (1) multiple imputation techniques such as multi-voxel spatial kernel imputation (Vaden et al., 2012), or (2) full information maximum likelihood estimation (Nelson et al., 2021). For instance, multiple imputation is known to provide variance that is most similar to estimates for voxels with no missing data. Future iterations of IBMMA will support a broader range of statistical designs, including longitudinal designs and survival analysis. Integrating machine learning frameworks could further increase IBMMA’s potential to identify biomarkers and improve the prediction of clinical outcomes.

The growing demand for processing and analyzing very large datasets of several thousand samples has revealed a significant disparity between the demand and the availability of relevant expertise. Currently, a small fraction of researchers possess the necessary training in data organization and analysis to meet these challenges (Schottdorf et al., 2024). Our efforts to close this gap led us to develop IBMMA by standardizing and simplifying statistical analysis of neuroimaging data. The IBMMA framework is designed for expansion into other domains that require sophisticated analysis of big data, offering a simple yet elegant approach to complex analytical tasks across various fields.

### Conclusion

IBMMA provides enhanced power and utility for analyzing large-scale, multi-site data by efficiently handling computational demands, managing missing data, and modeling site effects. As consortia efforts expand to tens of thousands of participants, IBMMA’s scalable architecture and flexible statistical modeling framework will be essential for reproducible, high fidelity brain mapping across diverse populations to accelerate discoveries in neuroscience and enhance neuroscience research.

## Acknowledgements

This study was supported by the Department of Defense award number W81XWH-12-2-0012; ENIGMA was also supported in part by NIH U54 EB020403 from the Big Data to Knowledge (BD2K) program, R56AG058854, R01MH116147, R01MH111671, and P41 EB015922. R01MH111671, R01MH117601, R01AG059874, MJFF 14848. The study was supported by ZonMw, the Netherlands organization for Health Research and Development (40-00812-98-10041), and by a grant from the Academic Medical Center Research Council (110614) both awarded to MO. The National Natural Science Foundation of China (No. U21A20364 and No. 31971020), the Key Project of the National Social Science Foundation of China (No. 20ZDA079), the Key Project of Research Base of Humanities and Social Sciences of Ministry of Education (No.16JJD190006), and the Scientific Foundation of Institute of Psychology, Chinese Academy of Sciences (No. E2CX4115CX). Funding from the SAMRC Unit on Risk & Resilience in Mental Disorders. R01MH105355-01A; NARSAD 27040; NIMH K01; MH118428-01. RO1 MH111671; VISN6 MIRECC. MH098212; MH071537; M01RR00039; UL1TR000454; HD071982; HD085850. Grant 01J05415 from the Special Research Fund (BOF) at Ghent University. German Research Foundation grant to J. K. Daniels (numbers DA 1222/4-1 and WA 1539/8-2). Supported by Czech Health Research Council AZV NU22-04-00661 and the core facility MAFIL supported by MEYS CR (LM2023050 Czech-BioImaging), part of the Euro-BioImaging (www.eurobioimaging.eu) ALM and Medical Imaging Node (Brno, CZ). Supported by a grant from the Ministry of Health of the Czech Republic, grant no. AZV NV18-7 04-00559. We acknowledge the core facility MAFIL supported by the Czech-BioImaging large RI project (LM2023050 funded by MEYS CR), part of the Euro-BioImaging (www.eurobioimaging.eu) ALM and Medical Imaging Node (Brno, CZ), for their support with obtaining scientific data presented in this paper. R21MH112956, R01MH119227, McLean Hospital Trauma Scholars Fund, Barlow Family Fund, Julia Kasaparian Fund for Neuroscience Research. K01MH118467; Julia Kasparian Fund for Neuroscience Research. R01MH113574. R01 MH106574. VA RR&D 1IK2RX000709. VA RR&D 1K1RX002325; 1K2RX002922. VA RR&D I01RX000622; CDMRP W81XWH-08–2–0038. German Research Society (Deutsche Forschungsgemeinschaft, DFG; SFB/TRR 58: C06, C07). The Natural Science Foundation of Jiangsu Province (No. BK20221554), and the Foundation for the Social Development Project of Jiangsu (No. BE2022705). 1R01MH110483 and 1R21 MH098198. PHRC, Fondation Pierre Deniker and SFR FED4226. Dana Foundation (to Dr. Nitschke); the University of Wisconsin Institute for Clinical and Translational Research (to Dr. Emma Seppala); a National Science Foundation Graduate Research Fellowship (to Dr. Grupe); the National Institute of Mental Health (NIMH) R01-MH043454 and T32-MH018931 (to Dr. Davidson); and a core grant to the Waisman Center from the National Institute of Child Health and Human Development (P30-HD003352). R01 AG050595, R01 AG022381. VA CSR&D 1IK2CX001680; VISN17 Center of Excellence Pilot funding. VA National Center for PTSD. The Beth K and Stuart Yudofsky Chair in the Neuropsychiatry of Military Post Traumatic Stress Syndrome.

