## Supplementary Materials for "Image-Based Meta- and Mega-Analysis (IBMMA): A Unified Framework for Large-Scale, Multi-Site, Neuroimaging Data Analysis"

| **Site** | ***N*** | **Age (mean)** | **Age (SD)** | **% Female** | **% PTSD** |
| --- | --- | --- | --- | --- | --- |
| adni_f1 | 74 | 65.83 | 1.63 | 0.00 | 58.11 |
| amc_f1 | 73 | 39.95 | 10.00 | 45.21 | 50.68 |
| beijing_f1 | 66 | 49.02 | 10.60 | 56.06 | 62.12 |
| cape_town_f1 | 62 | 27.55 | 5.73 | 100.00 | 1.61 |
| columbia_f1 | 59 | 36.14 | 12.70 | 42.37 | 37.29 |
| columbia_f2 | 18 | 36.56 | 12.44 | 55.56 | 11.11 |
| duke_f1 | 6 | 42.50 | 8.73 | 50.00 | 33.33 |
| duke_f2 | 24 | 41.08 | 11.22 | 25.00 | 33.33 |
| duke_f3 | 34 | 39.59 | 12.88 | 20.59 | 23.53 |
| duke_f4 | 49 | 41.90 | 9.29 | 12.24 | 30.61 |
| duke_f5 | 39 | 36.31 | 9.84 | 20.51 | 17.95 |
| emory_ed_f1 | 83 | 36.01 | 12.85 | 40.96 | 44.58 |
| emory_f1 | 51 | 39.92 | 12.18 | 100.00 | 27.45 |
| ghent_f1 | 54 | 36.23 | 11.48 | 100.00 | 11.11 |
| groningen_f1 | 41 | 38.37 | 9.65 | 100.00 | 100.00 |
| leiden_f1 | 3 | 19.67 | 0.58 | 100.00 | 66.67 |
| masaryk_f1 | 198 | 44.57 | 14.07 | 58.08 | 44.44 |
| mclean_kaufman_f1 | 49 | 36.14 | 12.82 | 100.00 | 100.00 |
| michigan_f1 | 58 | 30.81 | 7.85 | 0.00 | 70.69 |
| milwaukee_f1 | 70 | 32.77 | 10.35 | 47.14 | 27.14 |
| minn_va_f1 | 60 | 31.52 | 8.14 | 0.00 | 100.00 |
| minn_va_f2 | 97 | 32.56 | 7.58 | 3.09 | 27.84 |
| munster_f1 | 45 | 26.07 | 6.83 | 80.00 | 40.00 |
| nanjing_f1 | 143 | 57.26 | 5.90 | 53.15 | 35.66 |
| ontario_f1 | 169 | 38.75 | 12.37 | 64.50 | 73.37 |
| toledo_f1 | 49 | 37.37 | 11.69 | 38.78 | 30.61 |
| tours_f1 | 33 | 29.21 | 8.93 | 100.00 | 27.27 |
| tygerberg_f1 | 33 | 27.12 | 6.43 | 100.00 | 0.00 |
| umn_f1 | 55 | 42.40 | 9.92 | 9.09 | 20.00 |
| uw_cisler_f2 | 24 | 38.92 | 6.76 | 100.00 | 100.00 |
| uw_cisler_f3 | 43 | 30.91 | 8.56 | 100.00 | 100.00 |
| uw_grupe_f1 | 36 | 30.69 | 6.21 | 11.11 | 52.78 |
| vanderbilt_f1 | 50 | 31.34 | 4.63 | 18.00 | 30.00 |
| vetsa_f2 | 215 | 61.79 | 2.63 | 0.00 | 13.02 |
| waco_va_f1 | 36 | 41.69 | 11.15 | 11.11 | 58.33 |
| waco_va_f2 | 12 | 31.67 | 4.94 | 16.67 | 100.00 |
| waco_va_f3 | 14 | 38.79 | 8.89 | 0.00 | 100.00 |
| west_haven_va_f1 | 57 | 34.53 | 9.57 | 8.77 | 61.40 |
| **Total** | **2282** | **41.39** | **14.45** | **43.03** | **44.22** |

**Table S1: Demographic information by site.** For sites that collected data from multiple MRI scanners, each scanner was treated as a separate site (e.g., duke_f1 and duke_f2).

| **Site** | ***N*** | **Age (mean)** | **Age (SD)** | **% Female** | **% PTSD** |
| --- | --- | --- | --- | --- | --- |
| Duke | 34 | 39.85 | 12.75 | 17.65 | 26.47 |
| Minn VA | 90 | 32.57 | 7.54 | 2.22 | 24.44 |
| Munster | 44 | 26.23 | 6.83 | 79.55 | 36.36 |
| Ontario | 159 | 38.83 | 12.39 | 62.26 | 72.33 |
| UMN | 48 | 42.71 | 10.02 | 10.42 | 18.75 |
| UW Grupe | 22 | 30.41 | 6.11 | 13.64 | 50.00 |
| **Total** | **397** | **36.10** | **11.47** | **37.78** | **45.84** |

**Table S2. Demographic information by site for comparison subsample.**

**Simulation Dataset Comparing IBMMA and Complete-Case Analysis Performance**

**Simulation Design**

Simulated brain data was used to evaluate the performance of IBMMA’s method of handling missing data compared to the traditional complete-case analysis which only analyzes neuroimaging features with no missing data. Methods for creating the simulation dataset are described in the main report. In brief, we simulated 1,000 brains as 50×50×50 voxel arrays with values drawn from a standard normal distribution. A ground truth signal (Cohen’s *d* = 0.2) was embedded in a spherical region (251 voxels) to model a case-control design with 50% prevalence.

Missing data was introduced using a missing completely at random mechanism with a missingness rate of 0.01% per subject. This resulted in 113,138 out of 125,000 voxels (90.5%) having complete data from every subject.

We compared two analytical approaches. The IBMMA approach analyzed all 125,000 voxels using only subjects with non-missing data for each voxel, with a minimum threshold of 100 subjects required for analysis of a given voxel. In contrast, the complete-case approach analyzed only voxels with complete data across all 1,000 subjects. This approach simulated the traditional approach used by most neuroimaging software.

**Results**

***Detection Performance in True Signal Region***

Detection performance was evaluated within the 251-voxel ground truth signal region across multiple statistical thresholds. At the conventional threshold of t > 1.96 (approximately p < 0.05), IBMMA detected the signal in 249 of 251 voxels (99.2%), while complete-case analysis detected signal in 230 of 251 voxels (91.6%). This improved performance was maintained across increasingly stringent thresholds, with IBMMA showing advantages of 7.2% at t > 2.3, 7.2% at t > 3.1, and 5.2% at t > 5.0.

| **Threshold** | **IBMMA Detection** | **Complete-Case Detection** |
| --- | --- | --- |
| t > 1.96 | 249/251 (99.2%) | 230/251 (91.6%) |
| t > 2.3 | 247/251 (98.4%) | 229/251 (91.2%) |
| t > 3.1 | 241/251 (96.0%) | 223/251 (88.8%) |
| t > 5.0 | 191/251 (76.1%) | 178/251 (70.9%) |

**Table S3. Detection rates at various statistical thresholds**

***False Positive Rates***

False positive rates were assessed across the voxels that contained no true signal. Results are presented in **Table S4**. While IBMMA showed a modestly elevated FPR across all thresholds, the differences were consistent and relatively small, suggesting that the increased detection power did not come at the expense of substantial loss of specificity. This modest increase in false positives is addressed by adjusting for the larger number of multiple comparisons via multiple comparison correction.

| **Threshold** | **IBMMA FPR** | **Complete-Case FPR** |
| --- | --- | --- |
| t > 1.96 | 16.42% | 14.78% |
| t > 2.3 | 12.55% | 11.27% |
| t > 3.1 | 6.03% | 5.41% |
| t > 5.0 | 0.62% | 0.57% |

**Table S4. False positive rates (FPR) at various statistical thresholds**

**Overall Performance**

Receiver operating characteristic (ROC) analysis revealed superior overall performance for IBMMA, with an area under the curve (AUC) of 0.985 compared to 0.941 for complete-case analysis, representing a 4.68% improvement. When examining sensitivity at a fixed 5% FPR, IBMMA achieved 92.0% sensitivity compared to 85.3% for complete-case analysis, corresponding to a 6.7% improvement in detection capability.

The observed ~5-7% improvement in detection sensitivity has important implications for neuroimaging research, where effect sizes are typically small and statistical power is often limited. In clinical neuroimaging studies investigating subtle brain changes associated with psychiatric or neurological conditions, this improved sensitivity could translate to identification of brain regions that would be missed by traditional complete-case approach
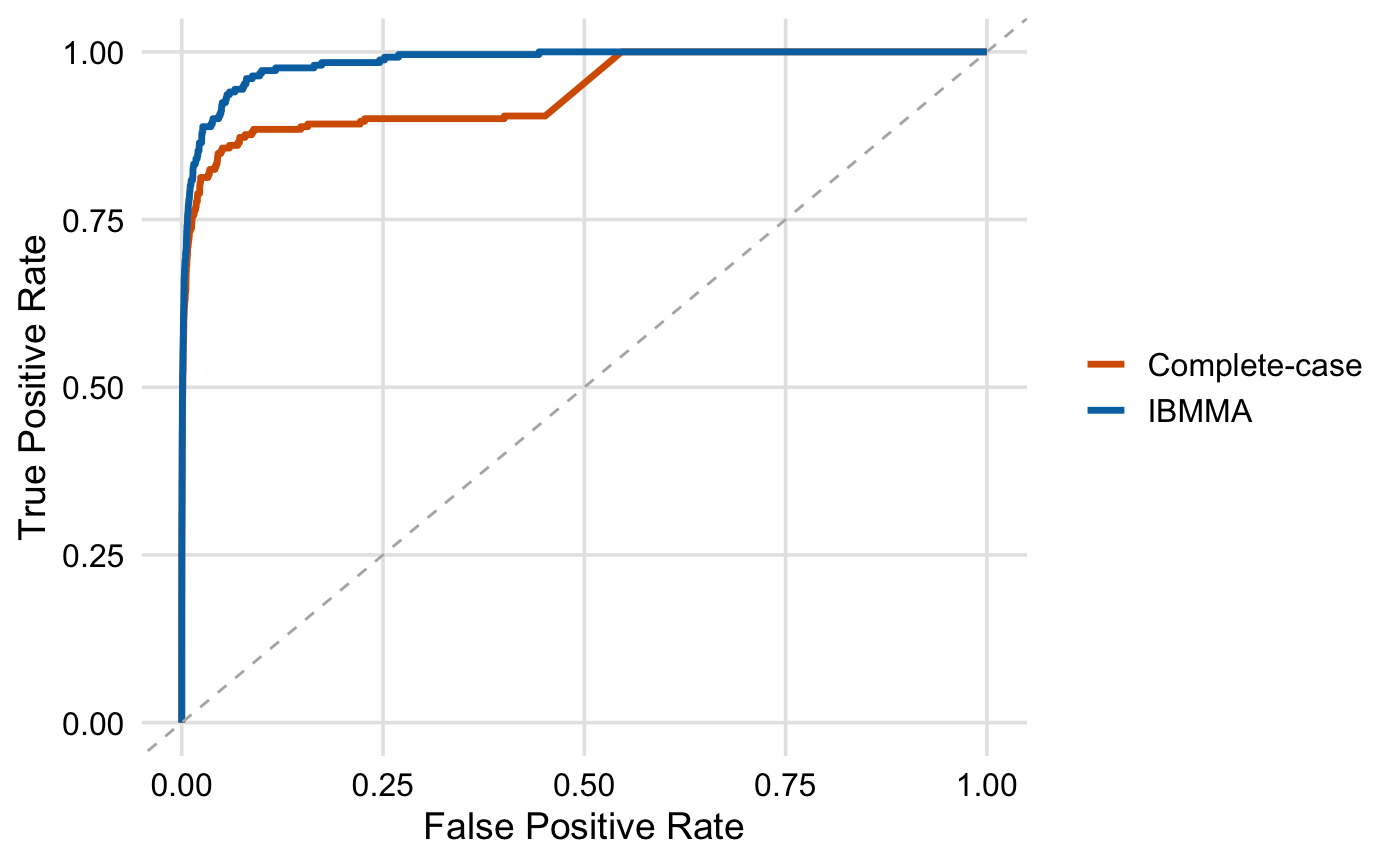
**Figure S1. Receiver operating characteristic analysis.** *ROC curves compare the detection performance of IBMMA-style (dynamic, voxelwise regression with partial missing data) and complete-case (analysis restricted to voxels with no missing data) methods in identifying true signal voxels. The ground-truth signal was simulated as a positive effect (cases > controls) within a spherical region. The IBMMA method yielded a higher area under the curve (AUC), indicating better sensitivity and specificity.*
